# Single-cell resolution spatial transcriptomic signature of the retrosplenial cortex during memory consolidation

**DOI:** 10.1101/2025.03.12.642891

**Authors:** Savannah R. Bliese, Budhaditya Basu, Stacy E. Beyer, Snehajyoti Chatterjee

**Affiliations:** Department of Neuroscience and Pharmacology, Carver College of Medicine, University of Iowa, Iowa City, IA, USA 52245; Iowa Neuroscience Institute, University of Iowa, Iowa City, IA, USA 52245; Interdisciplinary Graduate Program in Neuroscience, University of Iowa, Iowa City, IA, USA 52245

**Keywords:** Retrosplenial cortex, spatial memory, memory consolidation, gene expression, spatial transcriptomics

## Abstract

The retrosplenial cortex (RSC) is a critical brain region activated during spatial memory tasks and plays an underlying role in long-term memory consolidation. The RSC comprises multiple cell types, including different classes of excitatory neurons across laminar layers. These layer-specific cells form the hub of neuronal connection between the RSC and other brain regions, including the hippocampus. Despite the established role of the RSC in spatial memory, the transcriptomic signature of the neuronal sub-types in the RSC during spatial memory consolidation remained elusive. Here we used both unbiased and targeted spatial transcriptomic approaches to illuminate the transcriptional signature of the RSC following a spatial memory task. We found that genes related to transcription regulation, protein folding, and mitogen-activated protein kinase pathways were upregulated in the RSC after spatial learning during an early time window of memory consolidation. Further, cell type and excitatory neuronal layer-specific changes in gene expression were resolved using Xenium spatial transcriptomics. The distinct signatures of memory-responsive genes were observed in excitatory neurons across the laminar layers of the RSC following learning. Finally, we observed that blocking RSC excitatory neurons during the early temporal window after learning using a chemogenetic approach impaired long-term spatial memory. Overall, our results uncover a molecular signature of the RSC after learning and demonstrate the role of RSC excitatory neurons during the early time points of memory consolidation. This study underscores the importance of the learning-induced transcriptional signature of the RSC in long-term spatial memory consolidation and reveals a cell-type specific signature of memory-responsive gene expression.

## Introduction

The retrosplenial cortex (RSC) is a neocortical structure that acts as an integration center between sensory, limbic, and higher-order cortical brain regions[1]. In humans, the RSC has been shown to be integral for topographical and episodic memories[1, 2]. The RSC has also been identified as a key structure for spatial[3–8] and contextual fear memories[9–11] in rodents. The RSC integrates spatial navigational information using place-like and head-direction cells[12], and damage to the RSC leads to spatial disorientation[2] and both retrograde and anterograde memory loss[13, 14]. Neuronal tracing studies have identified several major RSC connections to other brain regions[15–17]. Among these include the hippocampus and entorhinal cortex, both of which are involved in a spatial memory circuit, and the connectivity between the RSC and these regions indicates its essential role in spatial navigation and memory[18]. The direct inputs from dorsal hippocampal sub-regions such as the subiculum and CA1[19, 20] are mediated by both glutamatergic[21] and GABAergic[22, 23] neurons. Furthermore, glutamatergic neurotransmission in the dorsal hippocampus and RSC has been implicated in contextual memory consolidation[9, 10, 24–26]. Spatial gene expression changes in the dorsal hippocampus during an early time window after learning is critical for long-term memory consolidation[27, 28]. However, the spatial transcriptional signature of the RSC during this early time point of memory consolidation has not been clearly described.

Recent single-cell transcriptomic studies have identified multiple spatially restricted excitatory neuronal sub-types within the RSC, many of which are unique to this region in comparison to other cortical areas[29, 30]. The RSC is anatomically diverse, comprised of granular and agranular subregions, both of which receive unique inputs and project to a wide variety of brain regions[31, 32], as well as display differing roles in behavior[21, 23, 33, 34]. The excitatory neurons within the RSC laminar layers exhibit functional and behavioral differences; for example, ablation of neurons in layers 4 and 5 of granular RSC caused amnesia in rats[35]. But, given this diversity, it has been challenging to delineate specific roles for each excitatory neuronal sub type within the RSC laminae. Therefore, it is critical to investigate the molecular signature of layer-specific patterns of excitatory neurons in the RSC to determine which neuronal populations are critical for long-term memory.

The molecular mechanism underlying spatial memory consolidation involves precisely timed transcriptional events. Memory-encoding neuronal ensembles comprising cells exhibiting the induced expression of immediate early genes (IEGs), also known as engram cells, have been extensively studied in the forebrain[36, 37]. Studies have shown the upregulation of activity-induced genes such as *Arc* and *Fos* in the RSC after learning in a contextual fear conditioning task[38, 39]. Using *in vivo* IEG imaging, a recent study showed engram activation in the RSC during a spatial memory task[40]. The study also demonstrated that the stability of the RSC engrams recruited during a spatial navigation task is associated with memory performance, providing evidence for a relationship between spatial memory and RSC engram ensembles[40]. Bulk transcriptomic profiling of the RSC and hippocampus identified a common induction pattern of an engram-specific gene expression signature after spatial learning in adult mice[41]. A study on the expression of Fos, an engram marker, suggests a unique role the RSC plays in tracking an animal’s position in the environment[42]. However, such learning-induced gene expression in the RSC at single-cell resolution across excitatory neuronal layers has never been performed. Therefore, understanding the precise transcriptomic signature of IEG that are markers of engrams in the RSC will provide mechanistic insights into the role of the RSC in spatial memory consolidation.

Single nuclei RNA sequencing and spatial transcriptomics have been used to study RSC-specific cell type identity[29] and gene expression following nerve injury[43]. However, the literature lacks single-cell and spatial transcriptomic studies investigating the anatomically distinct RSC neurons following a learning experience. In this study, we used cutting-edge molecular approaches to delineate the transcriptional signatures within the RSC during spatial memory consolidation. First, we analyzed gene expression in the RSC from our previously published spatial transcriptomics dataset after spatial learning[27, 44]. Our previous work used a deep-learning computation tool to predict neuronal activity across brain regions[44]. Additionally, we investigated the hippocampal gene expression using spatial transcriptomics after learning[27]. Thus, in the current study, we investigated the unbiased transcriptional signature in the RSC after learning. Next, we used a newly developed spatial transcriptomics platform, Xenium, to determine learning-induced gene expression in the RSC at single-cell resolution. Lastly, using a chemogenetic approach, we demonstrate the importance of learning-induced activation of excitatory neurons in the RSC for long-term spatial memory consolidation. Together, our findings provide insights into the transcriptomic signature of the RSC at single-cell resolution during long-term memory consolidation.

## Results

### Learning in a spatial object recognition task induces immediate early gene expression changes in the RSC

RSC neuronal activation has been reported during tasks that utilize spatial locations and landmarks[45–48]. Interestingly, introducing an object to the environment alters the mean firing rate of RSC neurons[48]. Therefore, we first investigated the unbiased transcriptomic signature of the RSC following learning in a spatial object recognition (SOR) task from our previously published datasets (GSE223066 and GSE201610)[27, 44]. Using this dataset, we have previously demonstrated the spatial transcriptomic signature of the dorsal hippocampus one hour after training in the SOR task, however, the learning-induced gene expression signature in the RSC was not studied. To investigate the gene expression changes in the RSC after spatial learning, we performed differential expression of genes (DEG) analysis on the RSC region comparing learning with homecage controls, and this analysis revealed 64 upregulated and 13 downregulated genes (**Fig. 1a-b, and Supplemental table 1**). Several of the top significant genes identified were IEGs such as *Fos*, *Arc*, *Nr4a1*, *and Egr1* (**Fig. 1c**). These activity-induced genes have been well characterized for their roles in learning and memory in the hippocampus and are used as markers of neuronal activity and engram ensembles[27, 38]. We further validated the induction of a few IEGs after learning from an independent experiment using qPCR analysis from RSC tissue (**Fig. 1d**). Next, we performed a gene ontology (GO) enrichment analysis to identify the pathways most represented among the DEGs in the RSC following learning (**Supplemental table 2)**. The top enriched pathways in the RSC include RNA polymerase II-specific DNA binding transcription factor binding, ubiquitin-protein ligase biding and ubiquitin-like protein ligase binding, unfolded protein binding, protein chaperone, heat shock protein binding, MAP kinase activity, MAP kinase phosphatase activity, ATPase regulator activity and misfolded protein binding (**Fig. 1e**). Genes related to DNA binding transcription factor binding such as *Nr4a1*, *Nr4a2*, and *Nr4a3* are closely linked to long-term memory consolidation and mice expressing a dominant negative form of Nr4a that blocks the transactivation function of all the Nr4a family members in the excitatory neurons in forebrain exhibits long-term memory deficits in contextual fear conditioning[49] and SOR tasks[50].

**Figure 1.**
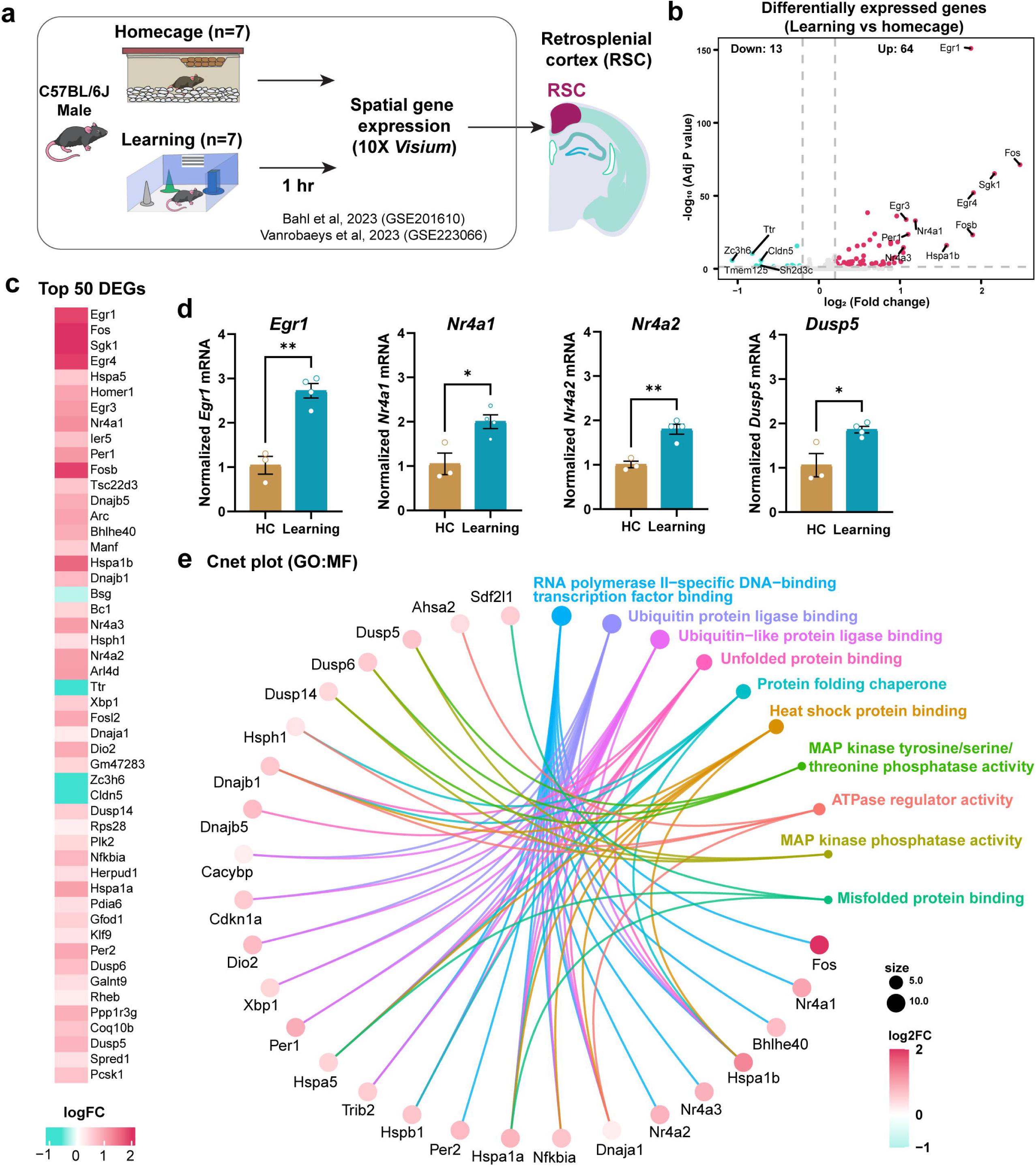
Spatial transcriptomic analysis using Visium reveals learning-induced gene expression in the RSC. **a** Schematics of the experimental paradigm being utilized for spatial gene expression and the region of interest being analyzed. **b** Number of differentially expressed genes found within the RSC 1 hour after learning (n=7 for both groups). **c** Heatmap displaying the top 50 significant differentially expressed genes within the RSC. **d** Expression of *Egr1*, *Nr4a1*, and *Dusp5* within the RSC 1 hour after learning. Normalized to homecage animals (homecage n=3, and SOR n=4). Unpaired t-test: *p < 0.05 (Nr4a1: p=0.0178, Dusp5: 0.0198), ** p < 0.005 (Egr1: p=0.0013). **e** Cnet plot exhibiting pathway enrichment analysis for the molecular function of the differentially expressed genes that were identified within the RSC.

### Single-cell resolution spatial transcriptomic analysis reveals distinct and overlapping learning-induced gene expression changes across major cell types in the RSC

Our unbiased gene expression analysis using the spatial transcriptomics (Visium) approach revealed a learning-induced gene expression signature in the RSC (**Fig. 1**). However, this approach lacks information regarding which cell types exhibit this learning-induced gene expression. Therefore, to determine the precise cell type-specific gene expression signature of the RSC, we utilized Xenium, an *in situ*-hybridization approach for single-cell analysis of each RNA molecule at a high spatial resolution within anatomically distinct tissue regions. We trained adult male mice in the SOR task and collected brains one hour after training for Xenium analysis. Homecage animals were used as baseline controls (**Fig. 2a**). We employed a 297-gene panel composed of a standard probe panel including cell-type markers, genes responsive to learning and memory, and genes involved in other processes (full probe list: **Supplemental table 3**). We obtained 26,484 high-quality cells from coronal sections in the RSC across learning and homecage animals with four biological replicates in each group.

**Figure 2.**
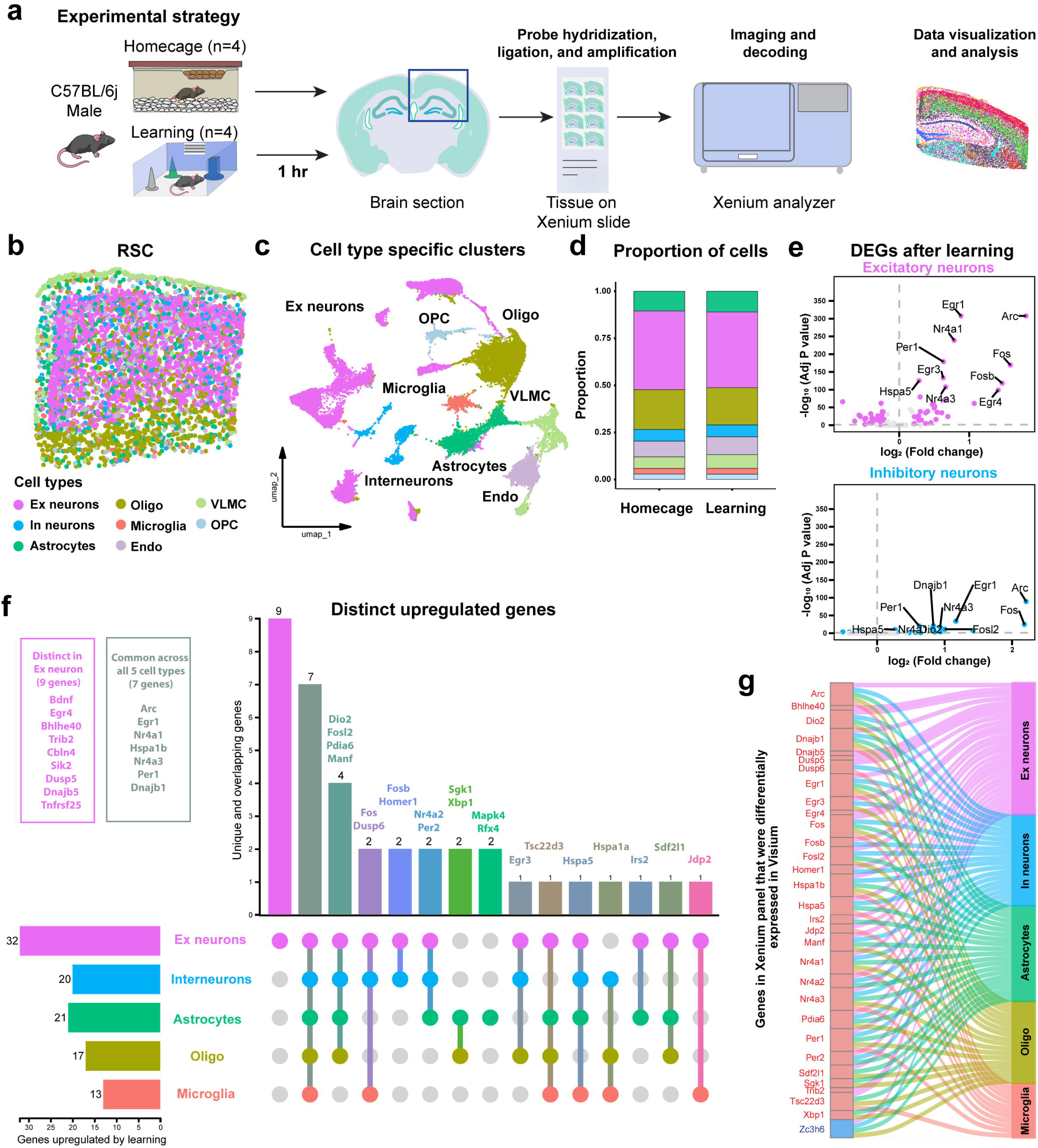
Single-cell resolution spatial transcriptomic approach reveals learning-induced gene expression across major cell types in RSC. **a** Schematics depicting the spatial learning task, the region of interest collected, and the processing for Xenium pipeline. **b** Post-processed image showing the alignment of annotated cell types in their correct target areas. **c** UMAP plot displaying unique cell types identified within the RSC. **d** Proportion plot comparing the proportion of cell types found among the two groups (homecage, n=4 and learning, n=4). **e** Volcano plots demonstrating the top significant differentially expressed genes for the excitatory (top panel) and inhibitory (bottom panel) neuronal cell types. Wilcoxon: FDR < 0.05, absolute log_2_ fold change > 0.2. **f** Upset plot illustrating the unique and overlapping DEGs across the major cell types identified within the RSC. **g** Sankey plot shows the common genes in both Xenium and Visium experiment that were significantly differentially expressed (red: upregulated, blue: downregulated) in the major cell types identified in Xenium analysis.

Within the RSC, we used the pre-defined marker gene expression in the Xenium gene panel to identify the different cell types. We identified eight major cell types within the RSC, including excitatory/glutamatergic neurons, inhibitory neurons, astrocytes, oligodendrocytes, endothelial cells, vascular leptomeningeal cells (VLMCs), microglia, and oligodendrocyte progenitor cells (OPCs) (**Fig. 2b-c**). The proportion of cells in each of the clusters between homecage and learning samples was comparable between the two groups (**Fig. 2d**). Given the role of neurons (excitatory and inhibitory) and glial cells (astrocytes[51], oligodendrocytes[52], and microglia[53]) in memory consolidation, we focused our learning-induced gene expression analysis on these five major cell types in the RSC. Among the 297 genes analyzed from the Xenium panel, a comparison between learning and homecage groups revealed 96 DEGs in excitatory neurons (32 up and 64 down), 23 DEGs in inhibitory neurons (20 up and 3 down), 43 DEG in astrocytes (21 up and 22 down), 44 DEG in oligodendrocytes (17 up and 27 down), and 14 DEG in microglia (13 up and 1 down) (**Fig. 2e, Supplementary** Fig. 1**, and Supplemental table 4**). Using an Upset plot, we examined the distinctly upregulated and downregulated genes after learning within these cell types in the RSC. Among the upregulated genes, *Arc*, *Egr1*, *Nr4a1*, *Hspa1b*, *Nr4a3*, *Per1,* and *Dnajb1* were induced after learning in all the five major cell types (excitatory and inhibitory neurons/interneurons, astrocytes, oligodendrocytes, and microglia), while *Bdnf*, *Egr4*, *Bhlhe40*, *Trib2*, *Cbln4*, *Sik2*, *Dusp5*, *Dnajb5*, and *Tnfrsf25* were induced exclusively in excitatory neurons. We also found that neuronal activity response genes *Fosb* and *Homer1* were exclusively upregulated in neurons, while *Sgk1* and *Xbp1* were exclusively upregulated in oligodendrocytes and astrocytes (**Fig. 2f**). Among the genes downregulated after learning, *Cirbp*, *Dner*, *Dpy19l1*, *Lypd6*, *Slc17a7* and *Garnl3* were downregulated in excitatory neurons, astrocytes, and oligodendrocytes. *Id2*, *Igfbp5*, *Parm1*, *Cdh20*, *Nrep*, and *Hpcal1* were downregulated in excitatory neurons, and oligodendrocytes. *Rbm3* was downregulated in excitatory neurons, interneurons, and oligodendrocytes. *Zc3h6* was downregulated in excitatory and inhibitory neurons, astrocytes, and oligodendrocytes, while *Plekha2* was downregulated in oligodendrocytes and microglia (**Supplementary** Fig. 2). Zinc finger CCCH containing 6 (Zc3h6) is predicted to have RNA binding and metal ion binding activity[54]. However, very little is known about Zc3h6 and its function in the brain. Comparing the learning-responsive DEGs from Visium and Xenium suggests that neurons in the RSC showed the highest overlap of learning-induced genes, particularly the upregulated genes in excitatory neurons (**Fig. 2g**). Therefore, our spatial transcriptomic analysis using Xenium revealed a cell-type specific signature of learning-induced genes in the RSC.

### Learning induces IEG expression changes across layer-specific neurons of the RSC

Our learning-induced gene expression analysis in the RSC using the Visium platform identified a higher number of upregulated genes in the RSC than downregulated genes. Importantly, we found a robust learning-induced upregulation of genes commonly used as engram markers in the RSC. RSC excitatory neuronal sub-types exhibit spatially recognizable laminar structural features[1, 29]. Additionally, these layer-specific glutamatergic neurons show unique electrophysiological characteristics[55, 56] and receive distinct inputs from the hippocampus[21, 23]. We performed unsupervised clustering from both the groups (homecage and learning) to identify the neuronal sub-types localized across different layers in the RSC. Investigating the expression of marker genes within each cluster of excitatory neurons further allowed us to classify the major layer-specific excitatory neuronal sub-types (**Supplementary** Fig. 3 **and Supplementary** Fig. 4). We identified seven major excitatory neuronal sub-types: layer 2/3 (L2/3), retrosplenial specific layer 2/3 (L2/3 RSP), layer 4 (L4), retrosplenial specific layer 4 (L4 RSP), layer 5 (L5), layer 6 (L6), and near projecting-subiculum NP SUB)) (**Fig. 3a-b**). Similarly, we also identified five major inhibitory neuronal populations (Pvalb, Sst, Vip, Lam5, and Sncg) along with glial cells (astrocytes, oligodendrocytes, microglia and OPC) in the RSC (**Fig. 3a-d**).

**Figure 3.**
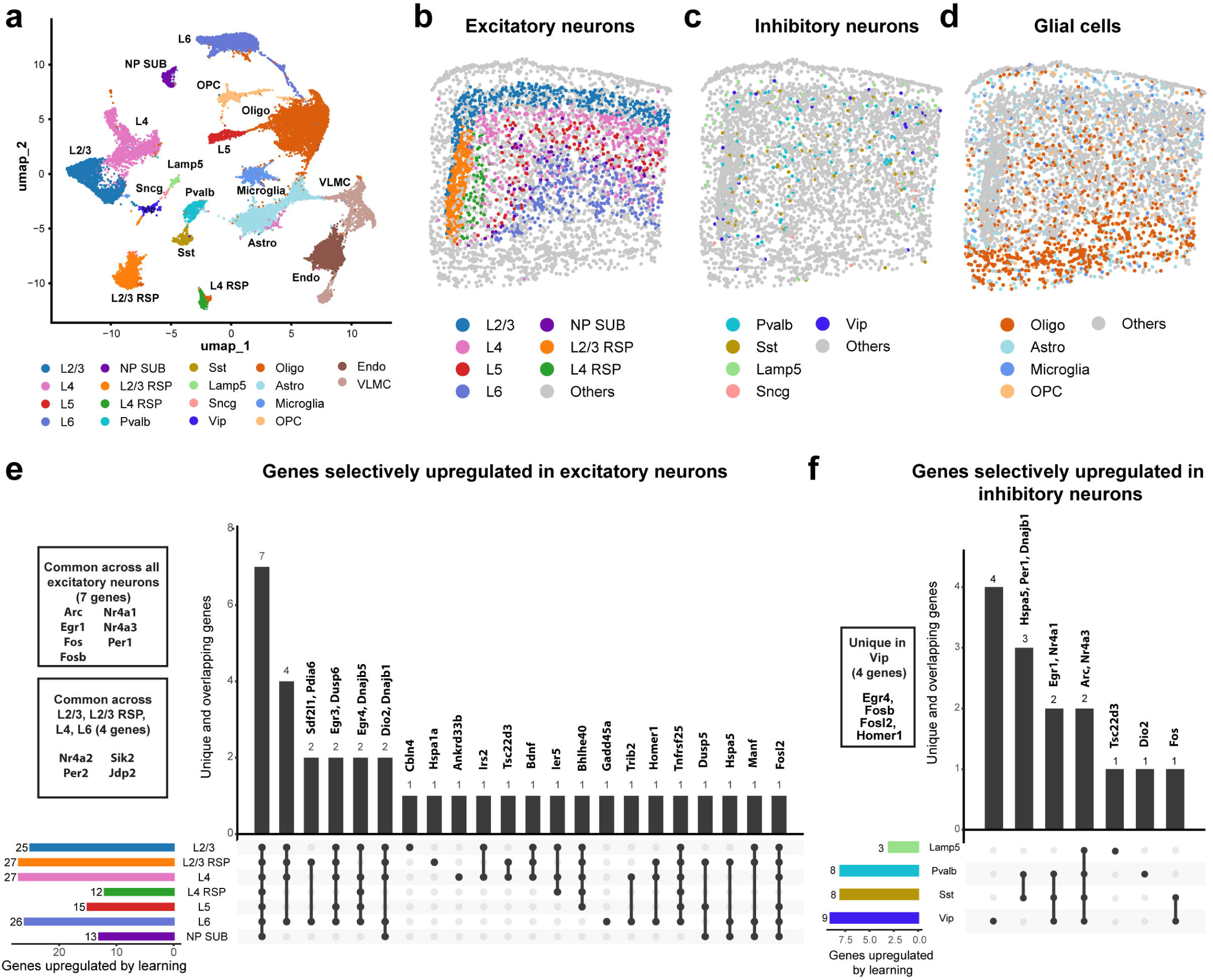
Single-cell resolution spatial transcriptomic analysis reveals the induction of IEGs after learning within different classes of RSC neurons. **a** UMAP exhibiting the layer-specific subclassification of cell types represented within major cluster identified within the RSC. **b-d** Spatial map of b) excitatory neuronal sub-types c) inhibitory neuronal sub-types and d) glial sub-types. **e-f** Upset plot showing the unique and overlapping expression patterns of upregulated genes across different e) excitatory neuronal sub-type and f) inhibitory neuron sub-types of the RSC.

To examine the learning-induced gene expression within the different populations of neurons, we performed DEG analysis on the 297 genes from the Xenium panel on all the neuronal sub-types (**Supplemental table 5)**. We used an Upset plot to investigate the learning-induced genes upregulated in each of the seven major excitatory neuronal populations identified in the RSC (**Fig. 3e**). Genes upregulated in only one sub-type of excitatory neurons include *Cbln4* upregulated in L2/3, *Hspa1a* in L2/3 RSP, *Ankrd33b* in L4, and *Gadd45a* in L6. Memory-related IEGs *Arc*, *Egr1*, *Fos*, *Fosb*, *Nr4a1*, *Nr4a3*, and *Per1* were commonly upregulated across all the seven major excitatory neuronal cell types, while TNF-receptor superfamily-related gene *Tnfrsf25* was upregulated in all the excitatory neuronal cell types except NP SUB. *Nr4a2*, *Per2*, *Sik2*, and *Jdp2* were upregulated in L2/3, L2/3 RSP, L4 and L6. Unfolded protein binding factors *Dio2* and *Dnajb1* were upregulated in L2/3, L2/3 RSP, L4, L6, and NP SUB, while chaperones *Sdf2l1* and *Pdia6* were upregulated in L2/3 RSP and L6. MAPK pathway-related genes *Egr3*, *Dusp6* were upregulated in L2/3, L2/3 RSP, L4, L5 and L6. Brain-derived neurotrophic factor and memory-related gene *Bdnf* was upregulated in L2/3, L2/3 RSP, and L4, and immediate early response gene *Ier5* was upregulated in L2/3, L4, and L4 RSP (**Fig. 3e, Supplemental table 5**). Among the inhibitory neurons, *Arc* and *Nr4a3* were commonly upregulated across all four inhibitory neuronal populations, while *Egr1* and *Nr4a1* were upregulated in Pvalb, Sst, and Vip interneurons. *Fos* was upregulated in Sst and Vip interneurons, and *Hspa5*, *Per1,* and *Dnajb1* were upregulated in Pvalb and Sst neurons (**Fig. 3f, Supplemental table 5**). Therefore, our spatial transcriptomic analysis after learning suggests an overall induction of IEG expression in the major neuronal populations, particularly across excitatory neuronal layers.

### Chemogenetic inhibition of the RSC excitatory neurons after learning impairs long-term spatial memory

Our spatial transcriptomics data identified robust learning-induced gene expression changes in the excitatory neurons of the RSC. To corroborate these findings, we used a chemogenetic approach to selectively manipulate the excitatory neurons immediately after learning, a critical time window for memory consolidation. We manipulated the activity of the excitatory neuronal population using a viral-based inhibitory designer receptor exclusively activated by designer drugs (DREADD) construct, which reduces the excitability of the neurons[57]. Expression of the construct was restricted to excitatory neurons within the RSC (**Fig. 4a**). Following training in the SOR task, animals were injected with clozapine-n-oxide (CNO, 2mg/kg). The virally expressed receptors respond specifically to CNO, allowing us to selectively block learning-induced activation of these neurons during the early time point immediately after learning. CNO treatment reduced the percentage of neurons expressing Fos, a marker of engram ensembles, within mCherry positive neurons (excitatory neurons infected by the DREADD virus) one hour after training in SOR, suggesting reduced neuronal activation in the CNO treated group (**Fig 4c-d**). Given the importance of engram ensembles in memory allocation during this period after learning[58], we aimed to test the long-term spatial memory when excitatory neurons within the RSC are selectively manipulated during this early time window. We found that mice injected with CNO immediately after training showed no preference towards the displaced object in SOR during the 24-hour long-term memory test (**Fig. 4e**). In contrast, mice injected with vehicle (saline) showed increased preference towards the displaced object during the test session compared to CNO injected mice (**Fig. 4e**). This finding suggests that reduced RSC excitatory neuronal activity during memory consolidation impairs long-term spatial memory (**Fig. 4e**).

**Figure 4.**
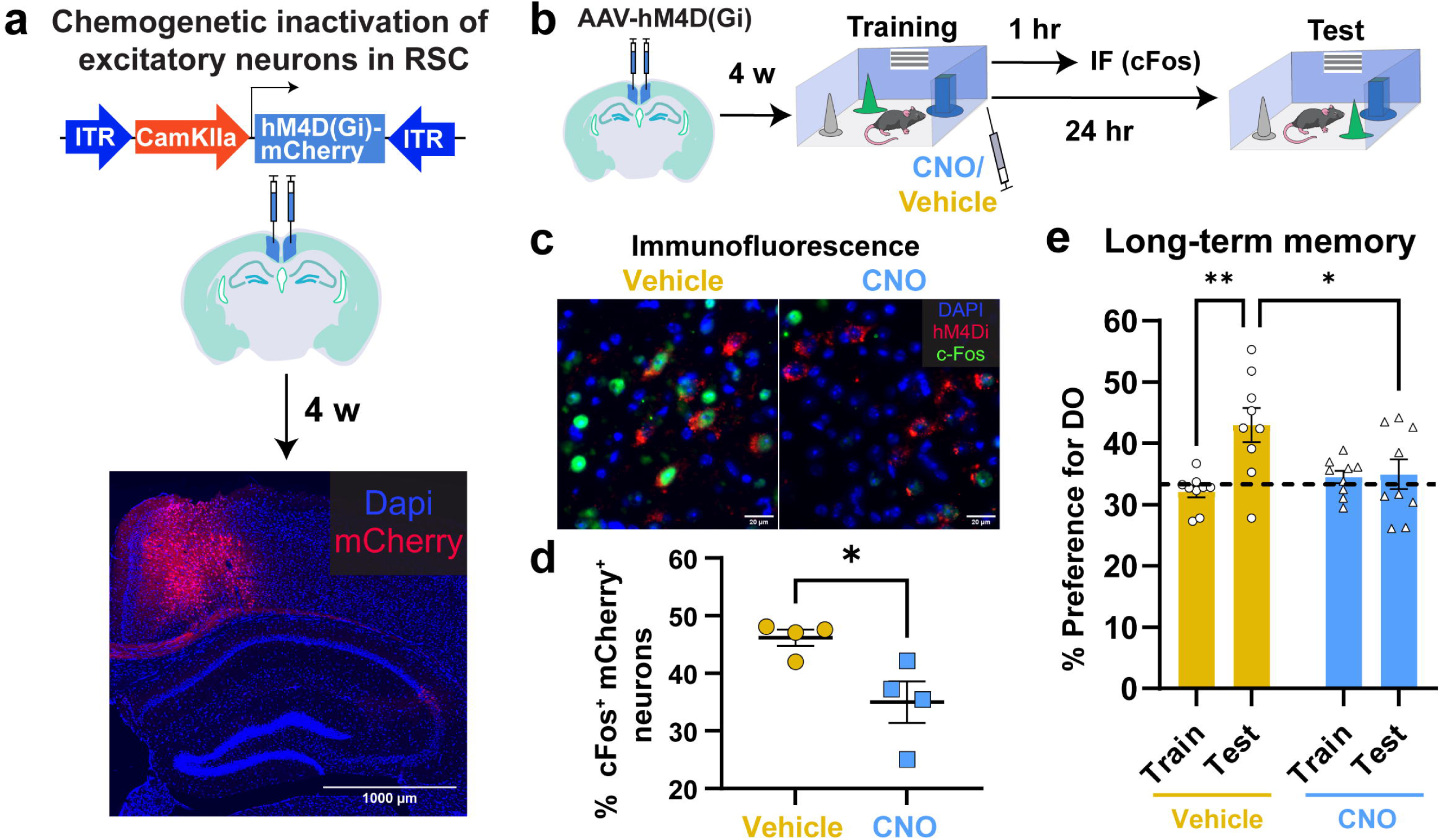
Activation of excitatory neurons after learning is essential for long-term spatial memory consolidation. **a** Graphic demonstration of the viral injection site, the viral construct that was used, and a representative immunofluorescence image showing the resulting viral expression within the RSC 4 weeks after infusion. **b** Schematic for the behavioral paradigm utilized. **c** Representative immunofluorescence images demonstrating Fos protein expression within DREADD positive neurons for the saline and CNO treated groups. **d** Dot plot showing the percent of DREADD positive neurons which are also positive for Fos protein expression 1 hour after learning on a SOR task. Unpaired T test: t (6) = 2.894, p = 0.0276. n=4 for both vehicle and CNO conditions. Error bars represent ± SEM. E. **e** Bar dot plot displaying behavioral performance on the SOR task between the saline and CNO injected animals. Long-term memory was assessed by finding the percent preference for the disturbed object (DO). 2 Way Anova: Significant trial type (Train-Test) x drug (vehicle-CNO) interaction: F (1, 32) = 6.856, p = 0.0134, main effect of trial type: F (1, 32) = 8.172, p = 0.0074. Šídák’s multiple comparisons test: vehicle: train vs test: p = 0.0030, test: vehicle vs CNO: p = 0.0429. n=9 for both vehicle and CNO conditions. Error bars represent ± SEM.

Overall, our results show the transcriptional signature of spatial memory in the excitatory neurons of the RSC and further demonstrate the importance of activation of excitatory neuronal ensembles in the consolidation of long-term spatial memory.

## Discussion

In this study, we uncover the spatial transcriptomic signature of the RSC during spatial memory consolidation. Using state-of-the-art spatial transcriptomic approaches, we demonstrate the cell type and spatially restricted excitatory neuronal layer-specific signature of the learning-induced genes in the RSC. Lastly, using a chemogenetic approach, we show that inhibiting the activation of RSC excitatory neurons after learning impairs long-term spatial memory. Our study provides molecular insights into the transcriptional signature of the RSC in long-term spatial memory and reveals the signature of engram-related genes during an early time window of memory consolidation.

Spatial transcriptomics provides the ability to investigate brain regions at high spatial resolution. We first utilized our previously reported spatial transcriptomic dataset[27, 44] to study learning-induced gene expression changes within the RSC. We found that pathways related to transcriptional regulation (RNA Pol II specific DNA binding factor binding), protein folding (unfolded protein binding, protein folding chaperone, heat shock protein binding, misfolded protein binding) and MAP kinase (MAP kinase tyrosine/serine/threonine phosphatase activity, and MAP kinase phosphatase activity) were enriched in the RSC following learning. These findings were complemented with spatial transcriptomics at single-cell resolution using the Xenium platform. We have previously shown that learning induces the expression of several genes linked to the engram ensemble in the hippocampal CA1 pyramidal region[27]. Using the same data set, we found similar learning-induced changes in engram-specific IEG expression in the RSC. Similar findings were reported previously by *Baumgärtel* et al., where they showed gene expression from the hippocampus and RSC following contextual fear conditioning and object exploration tasks[41], however, these studies lacked single cell resolution. They found an overlapping gene induction pattern following learning in the RSC and the hippocampus. Consistent with this study, our unbiased spatial gene expression analysis using the Visium platform within the RSC following SOR training identified similar upregulation of IEGs such as *Arc*, *Fos*, *Egr2*, *Fos*, *Arc*, *Per1*, *Nr4a1*, *Egr1*, and *Fosb*. Comparable upregulation of activity-induced genes such as *Arc*, *Fos, and Egr1* in the RSC after learning in a spatial memory[59] or a contextual fear conditioning task has been reported in other studies[38, 39]. Elucidating the functional impact of each upregulated gene in the RSC after learning is challenging. A recent study described the role of one key learning-induced gene, *Per1* in long term memory consolidation. We found that *Per1* was upregulated in all the five major cell types after SOR learning in the RSC. Interestingly, knockdown of *Per1* in RSC or hippocampal neurons was shown to impair long-term memory in a spatial object location task[60–62]. Furthermore, studies have shown that lesions or inactivation of the hippocampus impair activity-dependent expression of IEGs *Arc*, *Fos,* and *Egr1* in the RSC[63, 64]. This could point towards the complex entanglement between the hippocampus and the RSC, and this circuit’s role in learning-induced gene expression and memory consolidation.

The central purpose of our current study was to examine the cell type-specific transcriptomic signature of memory-related gene expression in the RSC. To our knowledge, this is one of the first studies to uncover transcriptional profiling of the RSC at spatial and single-cell resolution during the early time window of memory consolidation. Our cell-type-specific spatial gene expression analysis using the Xenium platform revealed learning-induced expression of engram-specific IEGs in the different neuronal populations of the RSC. We found that classical engram-specific memory responsive, IEGs such as *Arc*, *Egr1*, and *Nr4a1* were induced in all five major cell types (neurons and glial populations), while genes such as *Dusp5*, *Fosb*, *Homer1*, and *Egr4* were induced only within neurons, suggesting the importance of studying learning-induced gene expression at single cell resolution. Our targeted spatial gene expression experiment using the Xenium platform allowed us to examine genes that were predominantly induced in spatially restricted classes of neurons. Following training in the SOR task, we expanded our spatial transcriptomics work to examine the selective induction of memory-responsive gene expression across anatomically distinct excitatory neuronal layers in the RSC. These layer specific excitatory neurons play a critical role on RSC connectivity[32] and behavior[22, 34]. Although most of the classical IEGs considered markers of engrams were not restricted to any particular cell types or layers, we still found some regional and cell type-specific induction pattern of a few learning-induced genes. *Cbln4* was exclusively upregulated following learning in excitatory neurons in layer 2/3. Cbln4 is a secreted synaptic protein associated with inhibitory synapse recruitment[65]. One study showed a similar expression pattern of *Cbln4* in L2/3 of the RSC and found it was critical for synaptic density, which would have important implications on neuronal activity in the RSC[66]. We also found memory-related gene *Gadd45a* was upregulated exclusively in layer L6 excitatory neurons after learning. Gadd45a has previously been shown to have a role in spatial memory consolidation[67]—loss of Gadd45a impairs memory, while overexpression of Gadd45a in the hippocampus improves memory[68]. Therefore, our work provides the framework for future research to investigate the mechanistic underpinnings of these layer specific changes in learning-induced gene expression.

Our spatial transcriptomic analysis at single-cell resolution identified the upregulation of several memory-related genes in all excitatory neuronal populations in the RSC. Therefore, we used a chemogenic inactivation approach to determine the role of activation of excitatory neurons of the RSC after learning. Our inhibitory DREADD approach successfully blocked the learning-induced activation of Fos, a commonly used marker of neuronal activity and engram ensembles. We found that inhibiting the excitatory neurons in the RSC after learning impairs spatial memory. Thus, our work reveals that excitatory neuronal activation in the RSC during the early time window after learning is essential for long-term spatial memory consolidation. Previous inactivation studies have shown that the RSC and dorsal hippocampus are required for object location memory[69], with engrams being identified in these two brain structures[25, 40, 70]. However, the current theory of engrams in the field is that cortical and hippocampal engrams appear together, but those engrams must be matured to be engaged during retrieval[3, 40, 71, 72]. Interestingly, the RSC engrams generated after spatial learning are retained over prolonged periods, and the stability of those engrams during reactivation determines the strength of memory[40]. Thus, understanding the engram specific gene expression signature of the RSC during memory consolidation and its effect on behavior can provide us with valuable insights into neuronal dynamics that are instrumental in the storage of long-term spatial memory.

We utilized two cutting-edge spatial transcriptomics approaches to delineate the learning-induced gene expression signature of the RSC. The 55µm barcoded spot size of Visium provides a broad, unbiased transcriptomic analysis, while Xenium utilizes a targeted approach to investigate gene expression at single-cell resolution. The gene detection sensitivity of Visium is still lower than that of conventional transcriptomic approaches such as bulk RNA seq[73]. Thus, the low detection sensitivity of Visium could account for genes we identified to be induced by learning using Xenium but not Visium, such as *Bdnf* and *Gadd45a*. In conclusion, our study employed high-resolution spatial information to examine learning-induced genes in the RSC following SOR training. Even though our data is limited to a panel of genes, future work would investigate a larger pool of genes at single-cell resolution. Furthermore, the specific transcriptomic signature we identified across the laminar layers of the RSC suggests that each neuronal sub-type responds differently to activation related to memory consolidation. A recent study showed that IEG expression is reduced in layer 4 of the RSC in aged animals[74] which might provide the molecular basis of memory decline that is commonly seen with aging[75, 76].

Taken together, our work demonstrates a cell-type specific signature of learning-induced genes in RSC, laying the groundwork for future research to study the cell-type and layer-specific gene expression after learning in diseases associated with memory loss such as neurodegenerative diseases and age associated memory decline.

## Methods

### Data reporting

No statistical methods were used to predetermine sample size.

### Animals

Adult male C57BL/6J mice were purchased from Jackson Laboratories for all experiments. Mice had open access to food and water, and lights were maintained on a 12-hour light/dark cycle. All behavioral experiments were carried out in 3 to 4-month-old mice. Animals were randomly assigned to conditions when applicable. All experiments were conducted according to U.S. National Institutes of Health (NIH) guidelines for animal care and use and were approved by the Institutional Animal Care and Use Committee of the University of Iowa, Iowa.

### Drugs

Clozapine-n-oxide was dissolved in 0.9% sterile saline. For behavioral experiments, mice were injected intraperitoneally with a dose of 2.5 mg/kg immediately following training. 0.9% sterile saline was used as vehicle control.

### Adeno-Associated Viral (AAV) Constructs and Stereotaxic Surgeries

pAAV-CaMKIIa-hM4D(Gi)-mCherry was a gift from Bryan Roth (Addgene viral prep # 50477-AAV2). Mice were anesthetized with 5% isoflurane (Piramal Critical Care) for induction and then maintained at 2.0-2.5% during the surgical procedure. Betadine (Andwin Scientific) and ethanol pads were used to sterilize the scalp; then an incision was made down the midline to expose the skull. Holes were drilled into the skull above the RSC. The following coordinates (from Bregma) were used for viral injection into the RSC: 1.8 mm posterior, 0.45 mm lateral, and 0.75 mm ventral. A 33-gauge needle (WPI) attached to a 10 µl syringe was lowered into the RSC at a rate of 0.1 mm/10 sec. The needle rested at 0.65 mm below bregma for one minute then advanced down to 0.75 mm. Bilateral injections of the AAV constructs, 1 µl per side, were carried out at a rate of 100 nl/min, which was controlled by a micro syringe pump (UMP3; WPI). After the total volume was injected, the needle remained in place for an additional 2 minutes. The needle was raised slightly, then allowed to rest for an additional one minute before it was raised at a rate of 0.1 mm/15 sec. Bone wax (Surgical Specialties) was used to fill in the drill holes, and sutures were used to close the incision. Meloxicam at a dose of 5mg/kg was adminsitered as an analgesic and was given at the beginning of surgery and approximately 24 hours post-procedure.

### Spatial object recognition (SOR) task

Mice were single housed for seven days before the start of behavioral testing. Animals were handled for two minutes a day for five days total. Behavioral experiments were carried out from ZT0 to ZT3. Animals were brought into the room at least 15 minutes prior to experiment start. First, animals were allowed to explore the arena for 6 minutes for a habituation trial. Three objects were then placed into the arena and three training trials of six minutes each were carried out. 24 hours later, a single object was moved to a new location within the arena, and animals were tested for object location memory with a six-minute trial. Ethovision (Noldus) software was used to ensure arena settings remained the same between animals. Trials were manually scored and the amount of time the mouse investigated each object was recorded. To determine the percent preference for the displaced object, we took the total time spent exploring the moved object, divided by the total time exploring all objects, and multiplied that number by 100. For all spatial transcriptomics experiments, animals were still single housed for seven days and handled for five days. On the day of training, animals who were assigned to homecage condition remained in their cage, while animals assigned to the SOR condition carried out the habituation and training trials as described above. Whole brains or RSC tissue for these experiments were taken one hour after training, flash frozen, and then stored at −80C.

### Visium spatial transcriptomics data analysis

Count matrices from the previous published data (GSE223066 and GSE201610)[27, 44] were loaded into R (version 4.3.1) and a Seurat object was created using Load10X_Spatial function (Seurat package, version 5.1.0). Using Loupe Browser v8, we selected the Visium spots covering the RSC region and barcodes were exported as CSV files. The spatial barcodes were used to subset the RSC area from the whole coronal section. SCTransform normalization was performed on each replicate separately. Replicates from homecage controls and SOR samples were then integrated using a Seurat integration pipeline. Briefly, the integration anchors were found (FindIntegrationAnchors) from the list of 14 Seurat objects of homecage control and SOR samples. These anchors were used to integrate the 14 datasets together (IntegrateData). Principal component analysis (PCA) was performed using the RunPCA function (npcs=30). Based on 30 PCs, UMAP reduction was derived (using RunUMAP function). A k-nearest-neighbours graph was constructed based on Euclidean distance in PCA space and unsupervised Visium spots clustering was performed using a Louvain algorithm implemented in the Seurat package (FindNeighbors and FindClusters functions).

### Pseudobulk differential gene expression analysis

Instead of cluster based differential expression analysis, all Visium spots across the seven biological replicates from SOR group were pooled as pseudo bulk SOR group and spots from homecage (HC) controls across the seven biological replicates were pooled as pseudo bulk HC group. The pseudo bulk differential gene expression between SOR and HC was performed using the FindMarkers function with the following parameters: min.pct=0.2, test.use = “wilcox”. Each Visium spot was used as a replicate in this pseudo bulk analysis. The volcano plot and Sankey plot were generated using a custom ggplot2 script. Genes with an absolute log2 fold change > 0.2 and adjusted p value < 0.05 were considered significant.

### Molecular function enrichment analysis

The significant DEGs (p_val_adj < 0.05 & abs(avg_log2FC) > 0.2), including both upregulated and downregulated genes were further subjected to enrichment analysis executed using clusterProfiler v4.10. The Gene Ontology (Molecular Function) enrichment was performed using the enrichGO function implemented in clusterProfiler package with the following parameters: pvalueCutoff = 0.01, qvalueCutoff = 0.05, pAdjustMethod = “BH”, ont = “MF”. Further data visualizations were done using the clusterProfiler package.

### Xenium sample preparation and data acquisition

Brains were extracted from mice one hour after learning and then placed in −70° C isopentane for five minutes to freeze the tissue, then stored at −80^0^C. Mouse brains were embedded in OCT (Fisher Healthcare) and sectioned using a cryostat to 10 µm thick before being mounted on a Xenium slide (10x Genomics). Tissue was cut to mount only the RSC and hippocampus, which allowed for the placement of eight half-brain tissue samples per slide for further processing and analysis. All reagents used in Xenium sample preparation were purchased from 10x Genomics. Prior to probe hybridization, tissue was fixed in 4% paraformaldehyde for 30 minutes and then permeabilized using 1% SDS and chilled 70% methanol for 60 minutes. Slides were then placed into their cassettes and the pre-designed DNA probes were hybridized with the tissue overnight at 50°C. 297 DNA probes were pre-designed by 10x Genomics-247 included from their mouse brain gene expression panel, and 50 custom probes. After probe hybridization, samples were washed for 30 minutes followed by incubation with the ligation mix for two hours at 37°C. Immediately after, samples were washed then incubated for additional two hours at 30°C with the amplification master mix. Following another wash, samples were incubated overnight at 4°C in buffer before autofluorescence quenching and nuclei staining the following days. Slides were processed in the Xenium Analyzer. Briefly, fluorescent probes were hybridized to their targets, imaged, and then washed off to allow for subsequent probe cycles. Imaging detected puncta, which were decoded into gene ID’s and assigned a quality score. All data was acquired and further processed using R.

### Xenium analysis

#### Data preprocessing

The Xenium Ranger output files were imported into R (4.3.1) and a Seurat spatial transcriptomics object was created using the LoadXenium function (Seurat package v5.1.0). The data was filtered to remove cells with zero mRNA counts (using nCount_Xenium > 0). Subsequently, a region of interest (ROI) around the RSC was lassoed from the coronal section image (using Xenium explorer v3). This selected the RSC cells to be used for creating a Seurat object. The same process was followed for all biological replicates. The data from homecage control and SOR was normalized (using SCTransform) and integrated into a single Seurat object (using Seurat based FindIntegrationAnchors function). UMAP embeddings for the integrated object was computed using 30 principal components (RunPCA(npcs = 30), and RunUMAP(dims=1:30)). A k-nearest neighbor graph was constructed between every cell based on Euclidean distance in PCA space using FindNeighbors after which the unsupervised cell clustering was performed using the original Louvain algorithm (FindCluster (resolution=0.1)). We identified 16 clusters initially and further subclustered the inhibitory neuron clusters to increase the resolution. Cluster-specific markers were identified (using FindAllMarkers (logfc.threshold = 0.20)) and clusters were manually annotated from curated gene sets in the Xenium Panel. Two rounds of celltype annotations were performed. In the first round, major cell types identified were: excitatory neurons, inhibitory neurons, astrocytes, oligodendrocytes, microglia, endothelial cells, oligodendrocyte progenitor cells, and Vacuolar leptomeningeal cell (VLMC) were identified. In the second round, excitatory neurons were further classified into seven subtypes according to the cortical layers, inhibitory neurons were further clustered into five subtypes based on the markers *Pvalb, Sst, Lamp5, Sncg, Vip*. Cells were visualized using the Seurat based ImageDimPlot function.

#### Differential gene expression analysis

The differential gene expression analyses were performed between SOR and homecage control across different celltypes (using FindMarkers function), with the following parameters: min.pct=0.2, logfc.threshold=0, test.use = “wilcox”, assay = “SCT”. Genes with an absolute log2 fold change > 0.2 and adjusted p value < 0.05 were considered significant.

### Immunohistochemistry

Tissue for immunofluorescence staining and analysis was taken one hour after either the training or test session. Transcardial perfusions were performed using 4% paraformaldehyde (PFA, Sigma Aldrich) to fix brain tissue for preservation. Brians were removed and placed into 4% PFA for 24 hours. They were then placed in 10%, 20% and 30% sucrose solutions for 24 hours, or until the tissue had sunk. Tissue was sliced into 14-20 µm thick sections using a cryostat (Epredia). Slices were blocked with 5% normal goat serum (NGS, Jackson Immuno) for one hour before being incubated 24 hours at 4^0^C with a Fos antibody (SYSY Antibodies, 1:5000) diluted in 0.4% PBS with Triton-X at 4C. The following day slices were washed in 1x PBS three times then incubated in an AlexaFlour 647 secondary antibody (Jackson Immuno, 1:500) for two hours at room temperature (RT).

### Image Acquisition and Analysis

Stained tissue was mounted onto Superfrost Plus microscope slides (Fisherbrand) and coverslipped using Prolong Diamond Antifade Mounting Medium with DAPI (Invitrogen). Images of the RSC were taken with a 20x air objective using the VS200 Slide Scanner (Olympus). All images taken for Fos cell count used the same exposure time, laser, and gain settings. Images taken were imported into QuPath for cell count analysis. Briefly, region of interest was determined for the RSC and total cell count was found using the DAPI counterstain. Then the positive cell detection for mCherry and c-Fos were determined using predetermined thresholds. For c-Fos, the threshold was determined to be 200 percent of the mean fluorescence value of the background. Counts were then imported into Excel to find the percent the percent of cells expressing c-Fos for both mCherry+ cells and total cells, and the results were analyzed using Prism software.

### Total RNA extraction, cDNA preparation, and qPCR analysis

RSC tissue was extracted from animals one hour after learning on a spatial object recognition task and placed in RNAlater (Invitrogen) at −80° C. For RNA extraction, tissue was homogenized in Qiazol (Qiagen) with stainless steel beads (Qiagen), then added to chloroform and centrifuged at 4°C at 12,000g for 15 minutes. The aqueous solution was collected and added to EtOH, and subsequently washed using buffers and nuclease free water from the Qiagen RNAeasy Kit in a column tube (Qiagen). Samples were treated with DNase for 25 minutes at room temperature. Next, 100% EtOH, sodium acetate and glycogen were added to samples and allowed to rest for one hour at −20°C before being centrifuged at top speed for 20 minutes at RT. Supernatant was removed and 80% EtOH was added to the sample and spun again for five minutes before aspirating the remaining EtOH. Samples were allowed to dry before being resuspended in nuclease free water. RNA concentration was estimated using a Nanodrop (ThermoFisher), and estimates were used to produce a solution containing 1 µg of RNA. cDNA was prepared using the SuperScript™ IV First-Strand Synthesis System (Invitrogen) and diluted to 2 ng/µl. 2.25 µl of each sample was added to a 384 well plate along with 2.5 µl of Fast SYBR™ Green Master Mix (ThermoFisher) and 0.25 µl of a 5 µM primer mix (IDT). Real-time PCR was carried out using a QuantStudio 7 Flex Real-Time PCR System (Applied Biosystems, Life Technologies). Samples were run in triplicates and all data was normalized to housekeeping genes. The 2^(-ΔΔCt)^ method was used for gene expression analysis. The following sequences were used to create custom primers for use with qPCR: *Egr1* (FW: 5’ TGAACAACGAGAAGGTGCTG 3’; Rev: 5’ AGCGGCCAGTATAGGTGATG 3’), *Nr4a1* (FW: 5’ AAAATCCCTGGCTTCATTGAG 3’; Rev: 5’ TTTAGATCGGTATGCCAGGCG 3’), *Dusp5* (FW: 5’ GACAGCCACACTGCTGACAT 3’; Rev: 5’ AGGACCTTGCCTCCTTCTTC 3’), *Tubulin* (FW: 5’ ATGCGCGAGTGCATTTCAG 3’; Rev: 5’ CACCAATGGTCTTATCGCTGG 3’), *Pgk1* (FW: 5’ CGAGCCTCACTGTCCAAACT 3’; Rev: 5’ TCTGTGGCAGATTCACACCC 3’), *Actb* (FW: 5’ TCAACACCCCAGCCATGTAC 3’; Rev: 5’ CGGAGTCCATCACAATGCCT 3’).

### Statistics

No software was used to predetermine sample size. For behavioral data, all statistical analyses were performed in GraphPad Prism using unpaired two-tailed t-tests and two-way ANOVAs with repeated measures as the within-subject variable. Sidak’s multiple comparison tests were used for post hoc analyses where needed. Differences were considered statistically significant when plJ<lJ0.05.

## Ethics

The Institutional Care and Use Committee at the University of Iowa approved all procedures on mice in this study.

## Reporting summary

Further information on research design is available in the Nature Portfolio Reporting Summary linked to this article.

## Data availability

The Xenium data generated in this study have been deposited in the Zenodo repository (https://doi.org/10.5281/zenodo.14630054).

## Code availability

The code used for Xenium and Visium data analysis can be accessed through GitHub https://github.com/ChatterjeeEpigenetics/RSC_learning_2025.

## Acknowledgments

We thank the Iowa Neurobank Core and the Neural Circuits and Behavior Core at the Iowa Neuroscience Institute for using their facilities. We thank the University of Michigan Advanced Genomics Core for technical support on spatial transcriptomics using *10X Genomics Xenium*. We thank Dr. Li-Chun (Queena) Lin and Zachary Niemasz for their technical assistance. We thank Dr. Rainbo Hultman for providing valuable input on the manuscript.

## Author contributions

S.C. conceived the idea, designed the experiments, and supervised the work. S.C. and S.R.B. wrote the manuscript with input from all the authors. S.R.B. performed viral infusion, immunofluorescence, and qPCR analysis. S.R.B. and S.E.B. performed behavior, biochemical, and molecular biology experiments and analyzed and interpreted the data. B.B. performed all the bioinformatic analysis from spatial transcriptomics data.

## Ethics declarations

The authors declare that do not have any conflicting interests.

## Supplementary information

Supplementary Information includes

Supplementary Figure and Supplementary Figure Legend 1-4.

