## Supplemental information for "Single-cell resolution spatial transcriptomic signature of the retrosplenial cortex during memory consolidation"

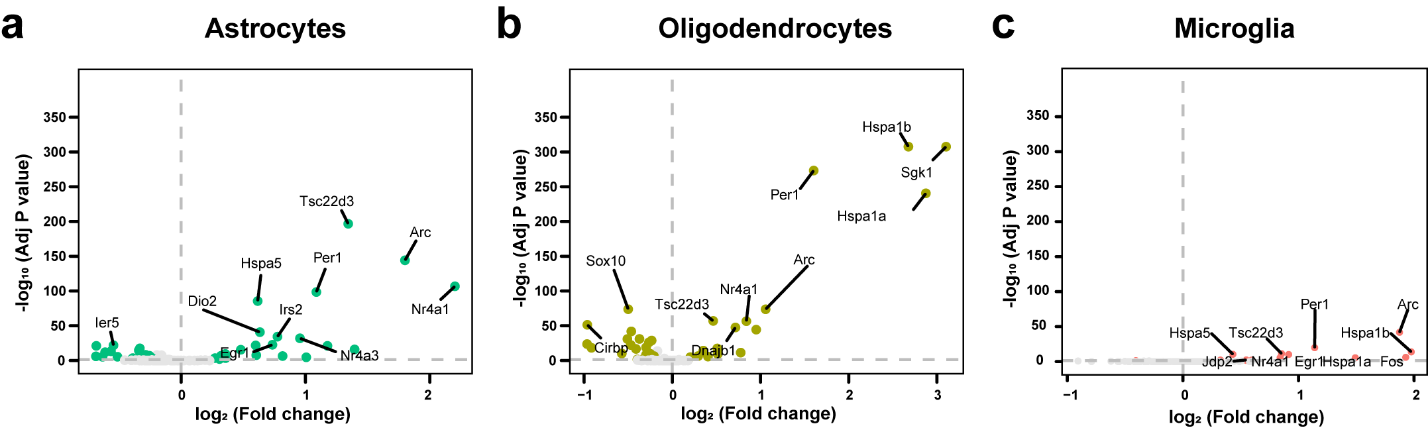

**Supplementary Figure 1. Learning-induced differential gene expression in non-neuronal cells.** Volcano plot showing significantly differentially expressed genes (FDR <0.05, log2 foldchange ±0.2 in **a** astrocytes, **b** oligodendrocytes, and **c** microglia.

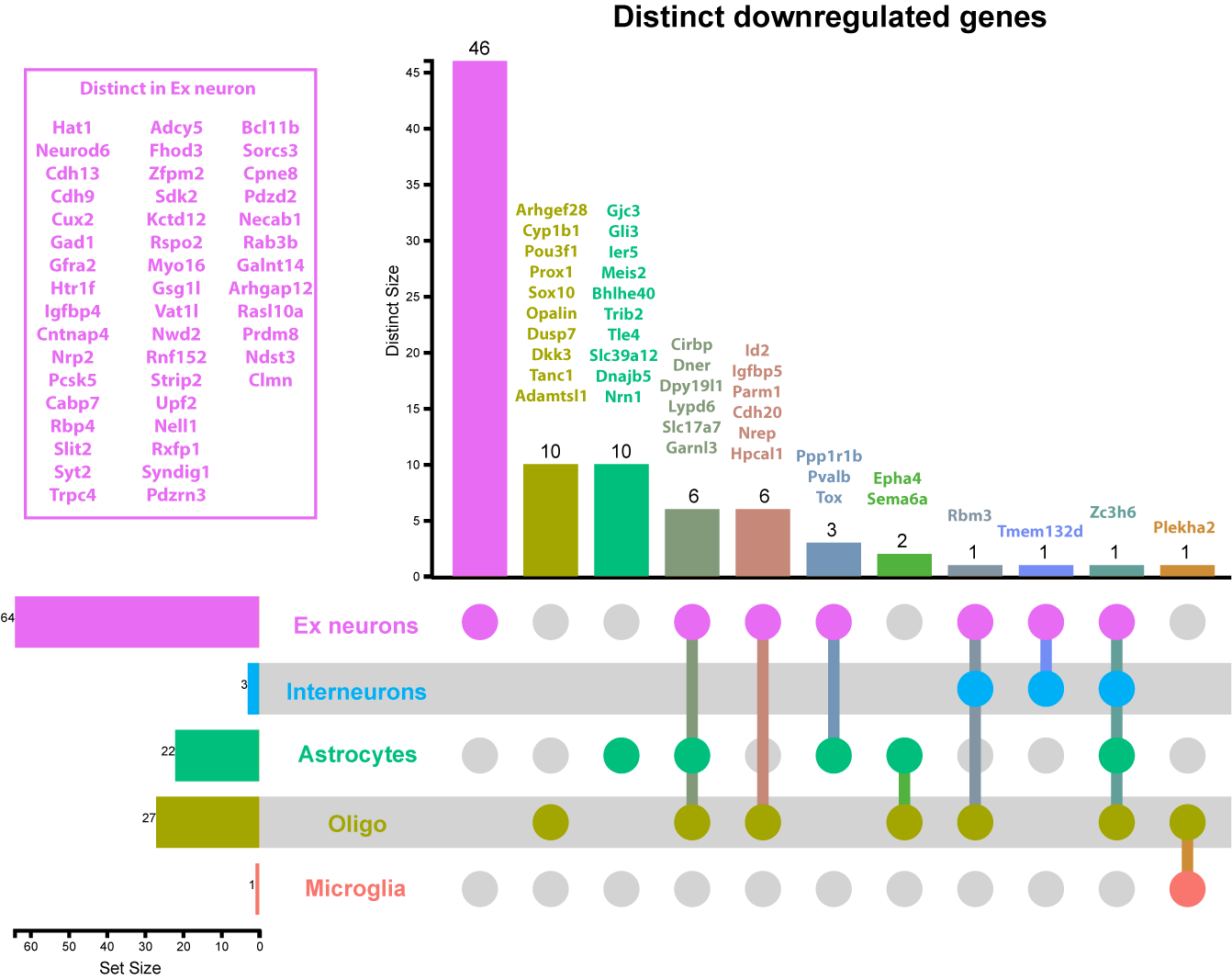

**Supplementary Figure 2.** **UpSet plot depicting all the significantly downregulated genes in the RSC across the five major cell types (excitatory neurons, inhibitory neurons, astrocytes, oligodendrocytes, and microglia) using the Xenium approach.**

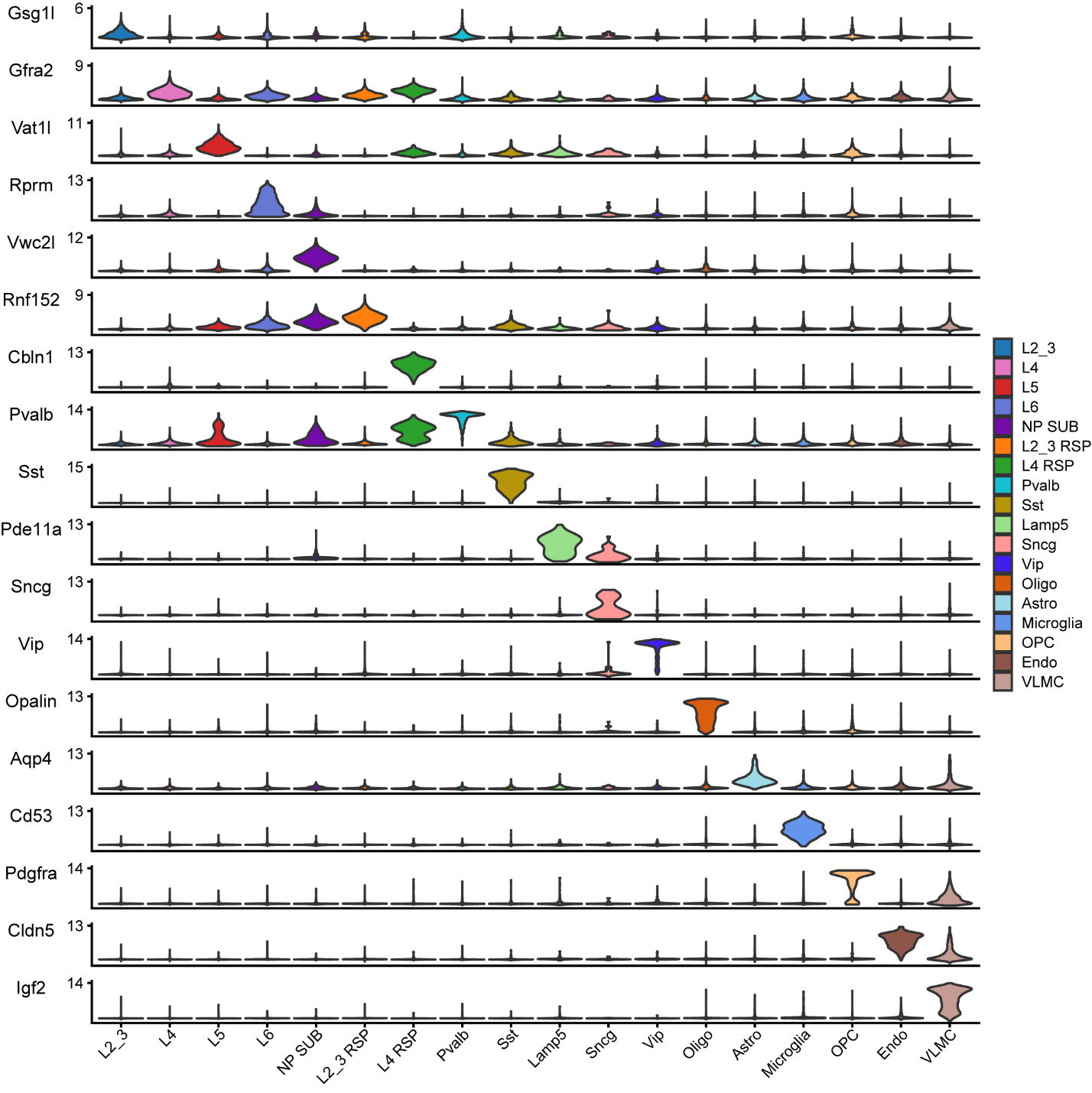

**Supplementary Figure 3.** **Violin plot depicting the expression of marker genes for each cell type.**

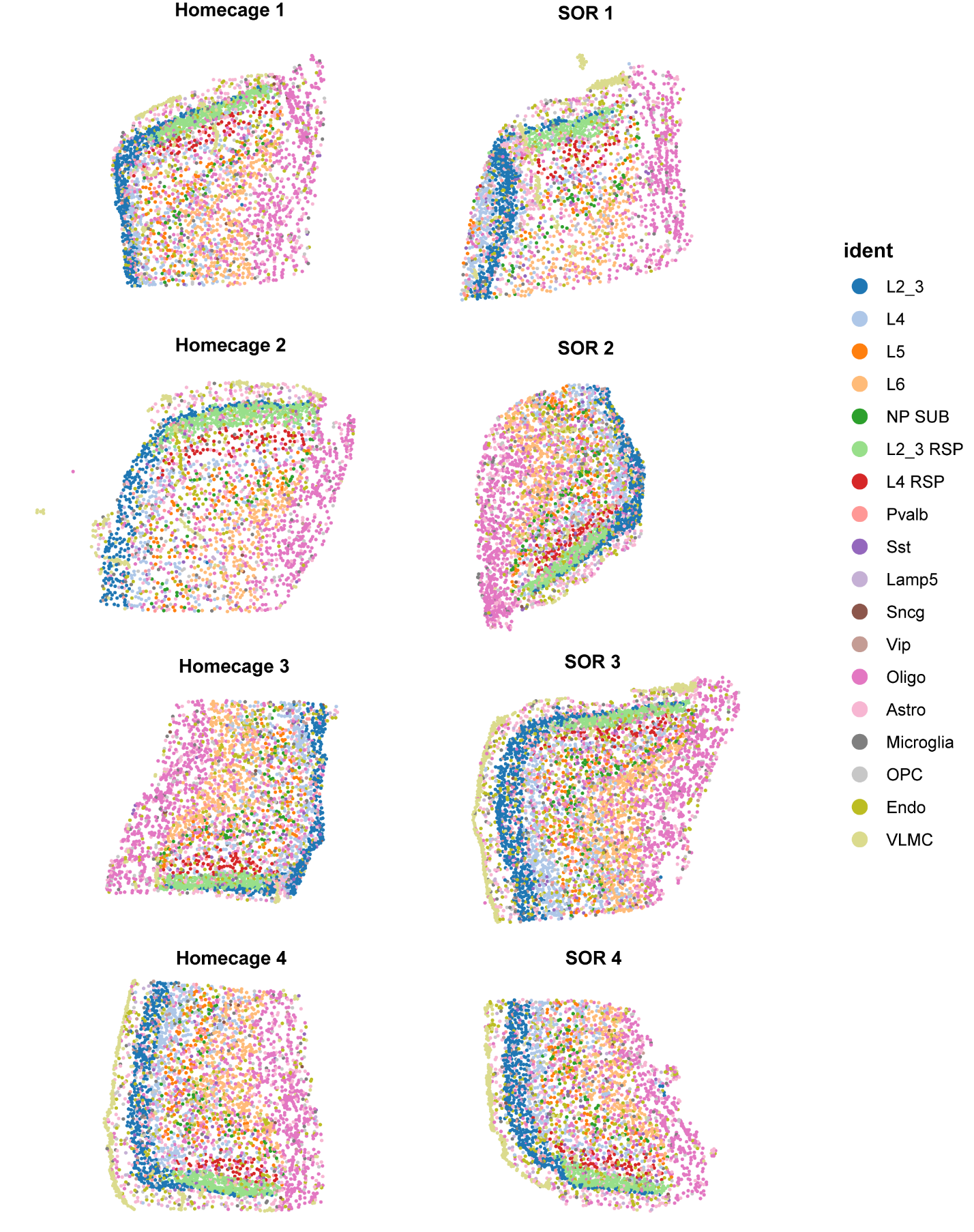

**Supplementary Figure 4.** Xenium sample replicates.
